# Invertebrate Metaxins 1 and 2: Widely Distributed Proteins Homologous to Vertebrate Metaxins Implicated in Protein Import into Mitochondria

**DOI:** 10.1101/2020.01.06.895979

**Authors:** Kenneth W. Adolph

## Abstract

Metaxin 1 and 2 genes, previously investigated in vertebrates, are shown to be widely distributed among invertebrates. But metaxin 3 is absent. The predicted proteins of the invertebrate metaxins were initially identified by homology with human metaxin 1 and 2 proteins, and by the presence of characteristic GST_Metaxin protein domains. Invertebrate metaxins were revealed for a variety of phyla, including Echinodermata, Cnidaria, Porifera, Chordata, Arthropoda, Mollusca, Brachiopoda, Placozoa, and Nematoda. Metaxins were also found in insects (Arthropoda) of different taxonomic orders: Diptera, Coleoptera, Lepidoptera, Hymenoptera, and Blattodea. Invertebrate and human metaxin 1 proteins have about 41% identical amino acids, while metaxin 2 proteins have about 49% identities. Invertebrate and vertebrate metaxins share the same characteristic protein domains, further strengthening the identification of the invertebrate proteins as metaxins. The domains are, for metaxin 1, GST_N_Metaxin1_like, GST_C_Metaxin1_3, and Tom37. For metaxin 2, they are GST_N_Metaxin2, GST_C_Metaxin2, and Tom37. Phylogenetic trees show that invertebrate metaxin 1 and metaxin 2 proteins are related, but form separate groups. The invertebrate proteins are also closely related to vertebrate metaxins, though forming separate clusters. These phylogenetic results indicate that all metaxins likely arose from a common ancestral sequence. The neighboring genes of the invertebrate metaxin 1 and 2 genes are largely different for different invertebrate species. This is unlike the situation with vertebrate metaxin genes, where, for example, the metaxin 1 gene is adjacent to the thrombospondin 3 gene. The dominant secondary structures predicted for the invertebrate metaxins are alpha-helical segments, with little beta-strand. The conserved pattern of helical segments is the same as that found for vertebrate metaxins 1, 2, and 3.

## 1. INTRODUCTION

The metaxins have previously been identified as vertebrate proteins involved in the import of newly synthesized proteins into mitochondria. Three vertebrate metaxins have been characterized: metaxin 1 (Bornstein et al., 1995), metaxin 2 (Armstrong et al., 1999), and metaxin 3 (Adolph, 2004, 2005). The vertebrate metaxins are thought to be components of protein complexes functionally equivalent to the TOM or SAM complexes of the outer mitochondrial membrane of yeast (*Saccharomyces cerevisiae*). The role of the TOM and SAM complexes in protein import into yeast mitochondria is much better understood than protein import in human cells and other vertebrate cells. This investigation concerns whether invertebrates, like vertebrates, have metaxin genes. The aims have been to understand which, if any, of the metaxins are found in invertebrates, how widely distributed they might be among invertebrate phyla, and, most importantly, fundamental properties of the metaxins encoded by invertebrate genomes. The results show that metaxin 1 and 2 genes are widely distributed among invertebrates, just as in vertebrates. But although metaxin 3 is highly conserved among vertebrates (Adolph, 2019), metaxin 3 was not found for the invertebrates studied.

The invertebrates in this investigation represent many of the invertebrate phyla. An emphasis was on invertebrates that are widely used as model organisms in biological and biomedical research. These include the classic model organisms *C. elegans* (Brenner, 1974; White et al., 1986) and *Drosophila melanogaster* (Wangler et al., 2017). Both had their genomes sequenced early in the history of whole genome sequencing (*C. elegans* sequencing consortium, 1998; Adams et al., 2000). Other organisms in this study that are important model systems and have had their genomes sequenced include *Ciona intestinalis* (Satoh, 2003; Dehal et al., 2002), *Nematostella vectensis* (Genikhovich and Technau, 2009; Putnam et al., 2007), and *Trichoplax adhaerens* (Schierwater and DeSalle, 2018; Srivastava et al., 2008).

The first metaxin gene to be identified was the mouse metaxin 1 gene (*Mtx1*) (Bornstein et al., 1995). The metaxin 1 protein was revealed to be located in the outer mitochondrial membrane, and experimental evidence demonstrated a role in protein import into mitochondria (Armstrong et al., 1997). The mouse metaxin 1 gene is in close proximity to the thrombospondin 3 gene (*Thbs3*) and the glucocerebrosidase gene (*Gba*). The thrombospondins are extracellular matrix proteins, and are important mediators of cell-matrix and cell-cell interactions (Adams and Lawler, 2011; Mosher and Adams, 2012).

A gene homologous to the mouse metaxin 1 gene was found to be present in humans at cytogenetic location 1q21. In human cells, the metaxin 1 gene (*MTX1*) is between a glucocerebrosidase pseudogene (*psGBA1*) and the gene for thrombospondin 3 (*THBS3*) (Long et al., 1996; Adolph et al., 1995), with a metaxin 1 pseudogene (*psMTX1*) also at this locus. The gene order is *GBA1*—*psMTX1*—*psGBA1*—*MTX1*—*THBS3*. In humans, a deficiency of the glucocerebrosidase enzyme as a result of mutations in *GBA1* can cause Gaucher disease (Mistry et al., 2017; Tayebi et al., 2003; LaMarca et al., 2004). An increased genetic risk for the development of Parkinson’s disease is also associated with *GBA1* mutations (Do et al., 2019).

A second metaxin protein was identified in the mouse through its interaction with metaxin 1 protein (Armstrong et al., 1999). The metaxin 2 protein shows 29% amino acid identities with mouse metaxin 1. The gene (*Mtx2*) is adjacent to a group of homeobox genes coding for transcription factors. Because of their interaction, metaxin 1 and metaxin 2 proteins might form a complex in the outer mitochondrial membrane involved in the import of proteins into mitochondria. A human gene (*MTX2*) homologous to *Mtx2* is at 2q31.2. Like the mouse gene, it is adjacent to a cluster of homeobox genes.

Zebrafish metaxin 3 was the first metaxin 3 to be identified (Adolph, 2004). The zebrafish metaxin 3 protein sequence, deduced from its cDNA sequence, was 40% homologous to zebrafish metaxin 1, and 26% homologous to metaxin 2. *Xenopus laevis* metaxin 3 was also identified through sequencing of cDNAs (Adolph, 2005). Zebrafish and *Xenopus* metaxin 3 proteins have 55% amino acid identities, indicating that the proteins are conserved to a great degree. The recognition that metaxin 3 is a metaxin but distinct from metaxins 1 and 2 was based on factors including amino acid sequence homology, presence of GST_Metaxin domains, phylogenetic analysis, and conserved patterns of protein secondary structure.

The import of proteins into mitochondria has been investigated in great detail in yeast (*Saccharomyces cerevisiae*) (Pfanner et al., 2019; Wiedemann and Pfanner, 2017; Neupert, 2015). In particular, the TOM complex (translocase of the outer mitochondrial membrane complex) and SAM complex (sorting and assembly machinery complex) bring about the import of beta-barrel proteins into yeast mitochondria and their insertion into the outer membrane. The Tom37 domain, a conserved protein domain of the SAM complex, is also present in the metaxin proteins, both vertebrate and invertebrate (this study).

Although the invertebrates include important research organisms used in many labs, nothing has been published about the metaxins of invertebrates. Metaxin proteins representing diverse invertebrate phyla have therefore been studied. The proteins were primarily those predicted by genome sequencing. Metaxin 1 and metaxin 2 were found to be highly conserved among invertebrates, and fundamental properties of the proteins were determined.

## 2. MATERIALS AND METHODS

The amino acid sequence alignments in Figure 1 were produced using the Global Align program of the NCBI (https://blast.ncbi.nlm.nih.gov/Blast.cgi; Needleman and Wunsch, 1970; Altschul et al., 1997). This program allowed the identities and similarities of two metaxin amino acid sequences to be determined along their entire lengths. The EMBOSS Needle program (https://www.ebi.ac.uk/Tools/psa/emboss_needle/) was also used to generate pairwise sequence alignments. The CD_Search tool (www.ncbi.nlm.nih.gov/Structure/cdd/wrpsb.cgi; Marchler-Bauer et al., 2017) was used to identify the protein domains included in Figure 2 through searching the conserved domain database of the NCBI. Evolutionary relationships of invertebrate metaxins (Figure 3) were examined with the COBALT multiple sequence alignment tool and the phylogenetic trees produced from the multiple sequence alignments (www.ncbi.nlm.nih.gov/tools/cobalt; Papadopoulos and Agarwala, 2007). In addition, the Clustal Omega program (https://www.ebi.ac.uk/Tools/msa/clustalo) was used to generate multiple sequence alignments and phylogenetic trees. The genes adjacent to invertebrate metaxin genes were identified from the related information on “Gene neighbors” that resulted from BLAST searches (https://blast.ncbi.nlm.nih.gov/Blast.cgi). Adjacent genes were also identified using the Genome Data Viewer and, earlier, the Map Viewer genome browsers (www.ncbi.nlm.nih.gov/genome/gdv). The PSIPRED secondary structure prediction server (bioinf.cs.ucl.ac.uk/psipred/; Jones, 1999) was used to predict the alpha-helical structures of invertebrate metaxin proteins in Figure 4. The presence of transmembrane helices was investigated with the PHOBIUS program (www.ebi.ac.uk/Tools/pfa/phobius/; Madeira et al., 2019) and also the TMHMM prediction server (www.cbs.dtu.dk/services/TMHMM/; Krogh et al., 2001).

**Figure 1.**
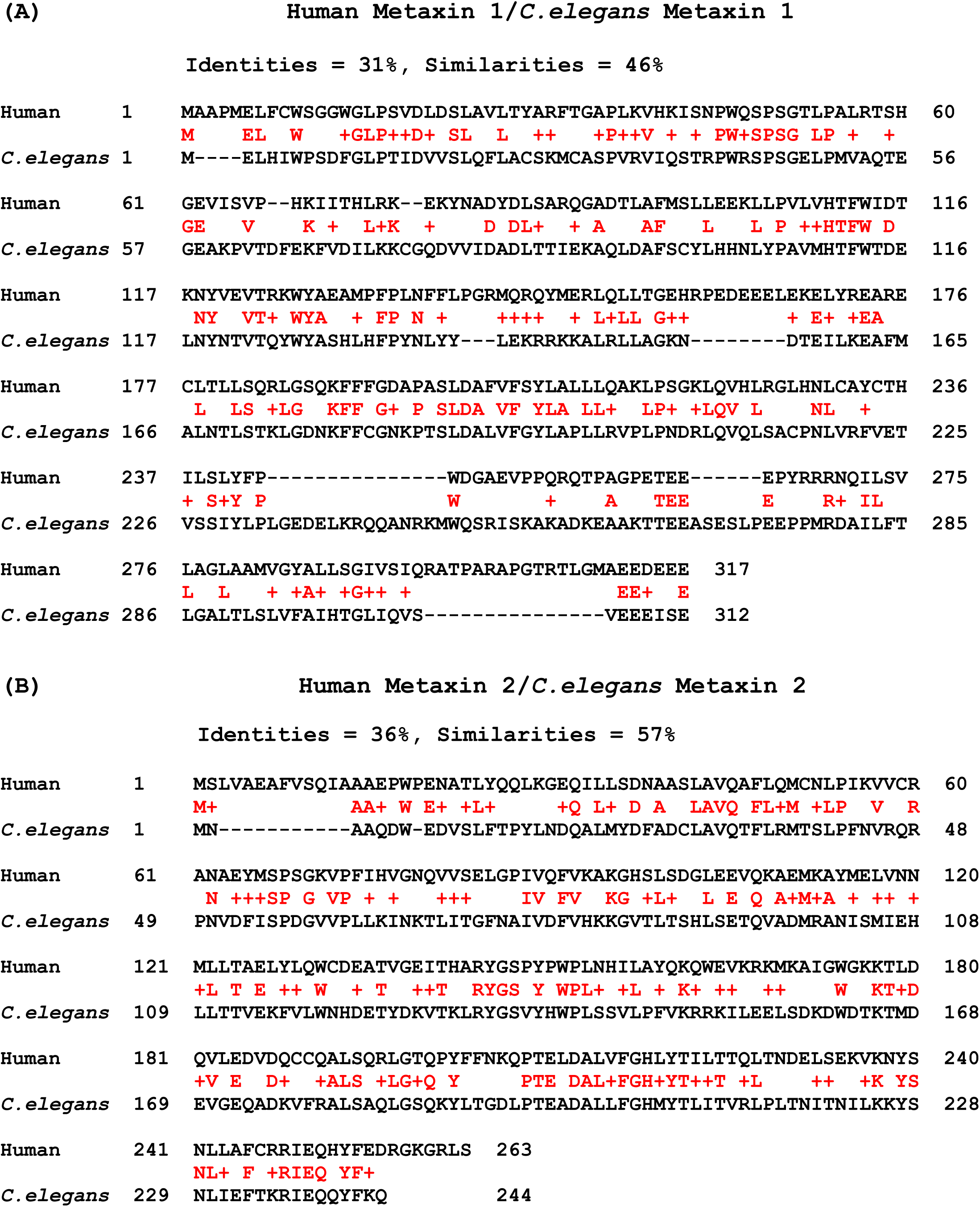
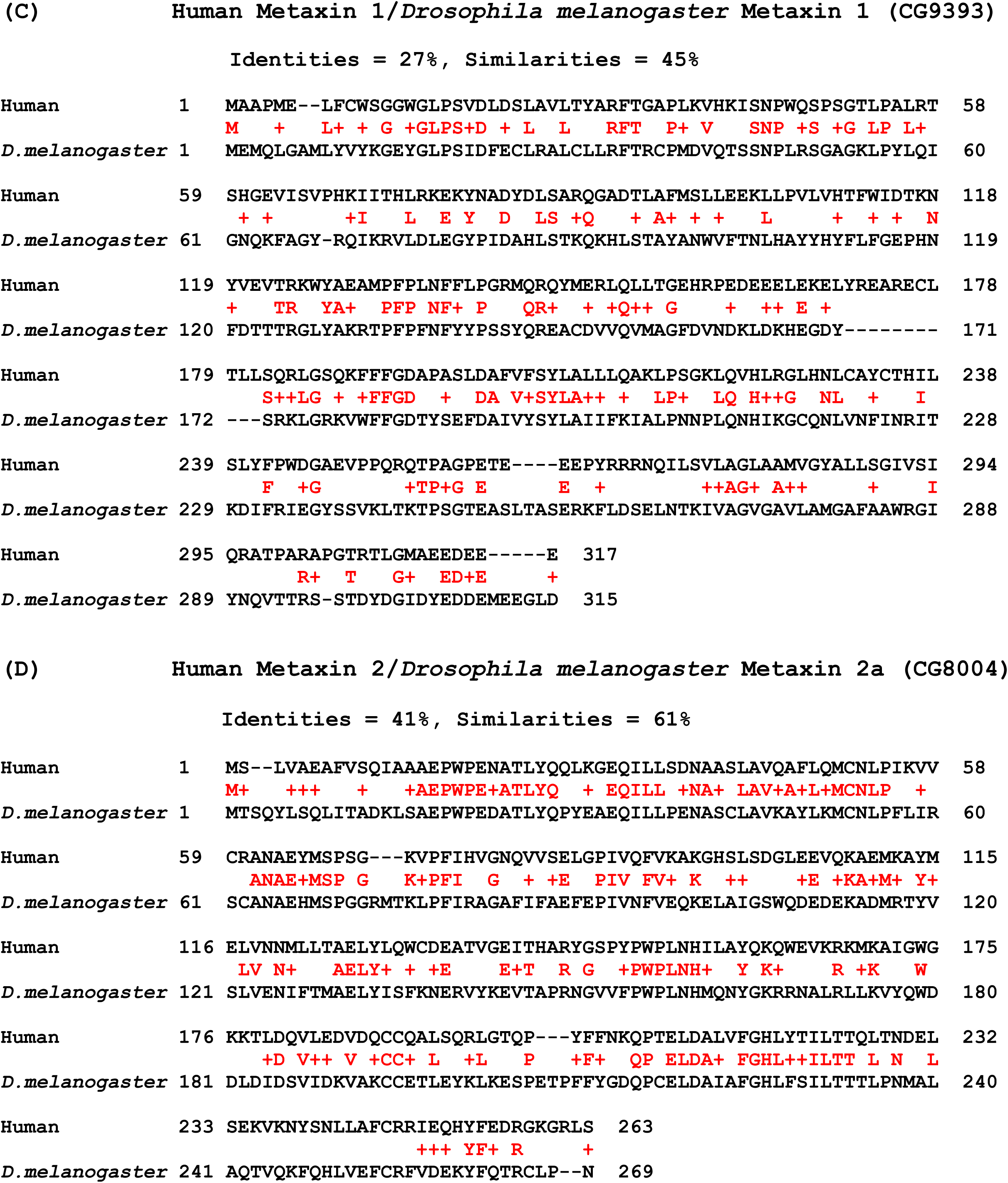
Homology of invertebrate metaxins: amino acid sequence alignments. (A) The amino acid sequence of human metaxin 1 (317 amino acids) is aligned with the sequence of a representative invertebrate *C. elegans* (312 aa), an important model system for studies of development. The identities and similarities are shown in red between the sequences. Identities are 31% and similarities are 46% for this example. Invertebrates, generally, share about 41% amino acid sequence identities with human metaxin 1. (B) The alignment of human metaxin 2 (263 aa) with *C. elegans* metaxin 2 (244 aa) shows 36% identical amino acids, with 57% similarities. Invertebrate metaxin 2 sequences more typically share about 49% identities relative to human metaxin 2. Comparing the sequences of various invertebrate metaxin 1 proteins with invertebrate metaxin 2 proteins shows about 27% identities. (C) Alignment of human metaxin 1 and an insect metaxin 1. In this alignment with a *Drosophila melanogaster* metaxin 1 homolog (315 aa), 27% identical amino acids and 45% similarities are found. *D. melanogaster* was included because it is a classic model system of genetics and developmental biology. Comparing the metaxin 1 sequences of insects of different orders with each other shows about 38% identities. (D) For human metaxin 2 and *D. melanogaster* metaxin 2a (269 aa), there are 41% identities and 61% similarities. *D. melanogaster* is one of the *Drosophila* species with two metaxin 2 proteins, 2a and 2b. The two proteins have 48% sequence identities. Comparing the metaxin 2 sequences of a variety of insects of different orders shows about 51% homology.

**Figure 2.**
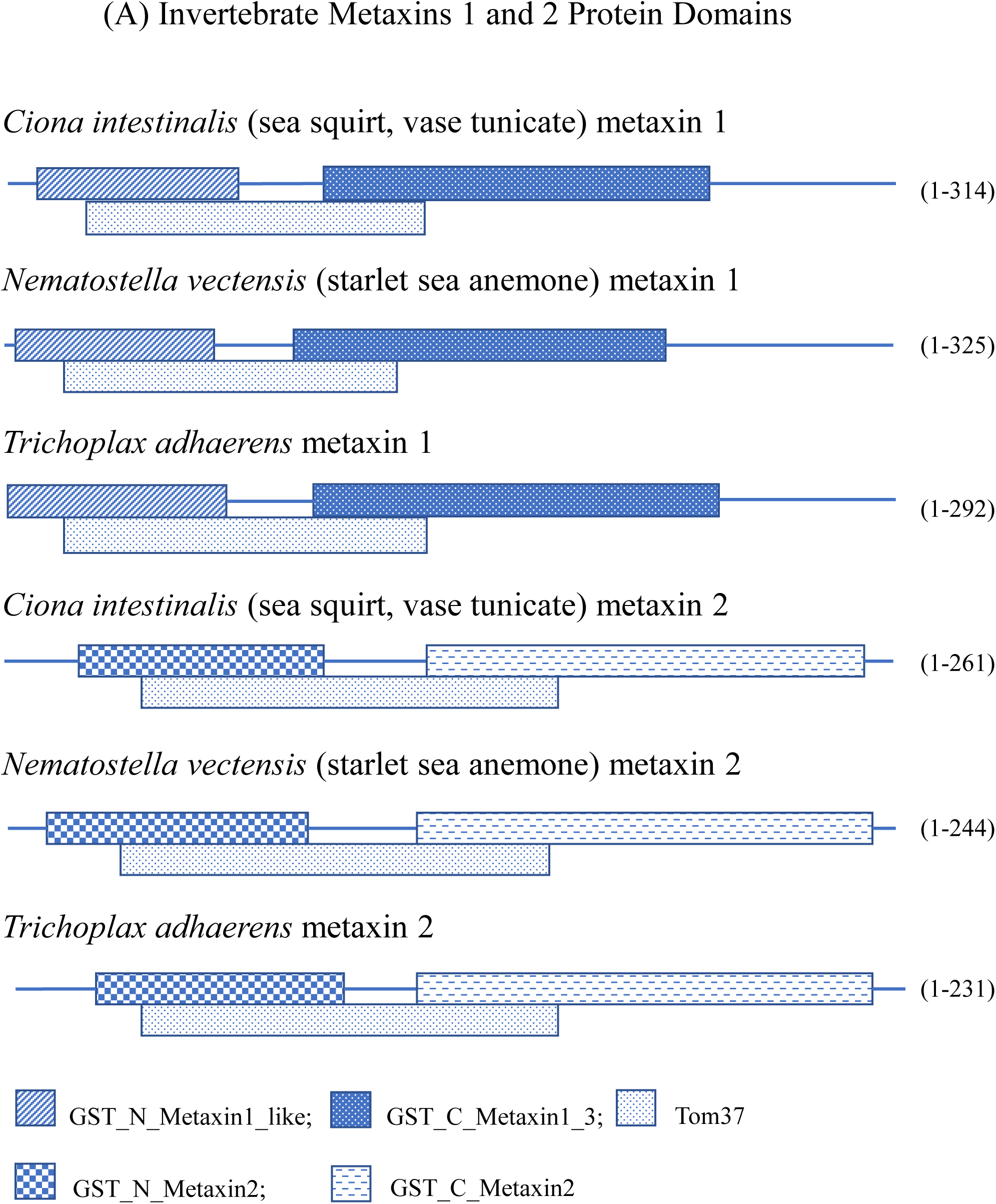
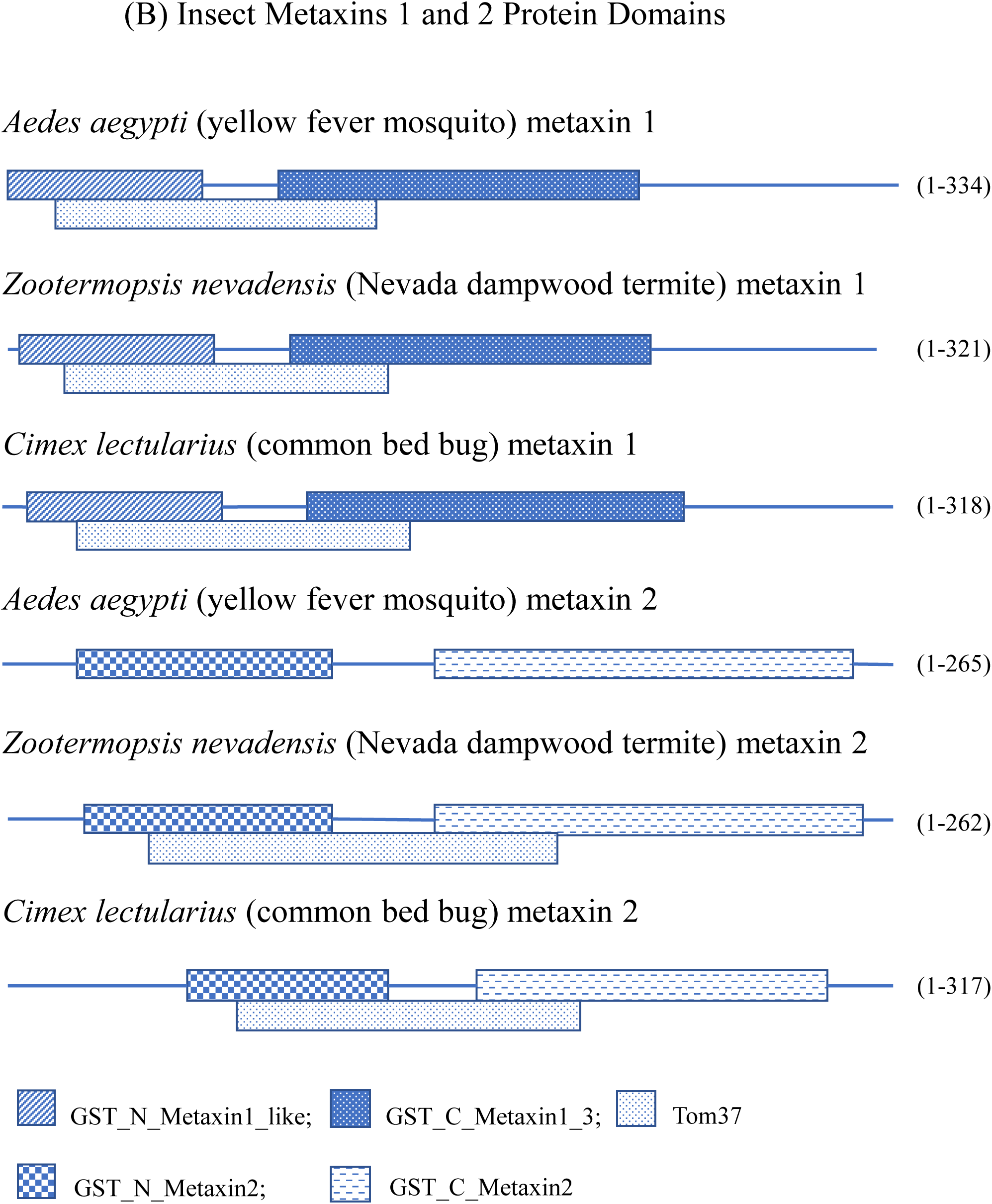
Domain structure of invertebrate metaxin proteins. (A) The metaxin 1 and metaxin 2 protein domains of representative invertebrates, not including insects, are shown. These are the characteristic GST_N_Metaxin, GST_C_Metaxin, and Tom37 domains. The invertebrate examples include a sea squirt (*Ciona intestinalis*; Phylum: Chordata), a sea anemone (*Nematostella vectensis*; Phylum: Cnidaria), and *Tricoplax adhaerens* (Phylum: Placozoa). The domain structures are highly conserved in these examples, and in the metaxins of other invertebrates. (B) The metaxin protein domains are shown for three representative insects: the yellow fever mosquito (*Aedes aegypti*; Order: Diptera), a termite (*Zootermopsis nevadensis*; Order: Blattodea), and the common bed bug (*Cimex lectularius*; Order: Hemiptera). Other insects have the same domains, and the insect domains are the same as for the invertebrates in (A). The presence of a Tom37 domain in both metaxin 1 and 2 proteins is in keeping with a possible role for the invertebrate metaxins in the import of proteins into mitochondria. Tom37 is a yeast protein with a well-characterized role in protein import.

**Figure 3.**
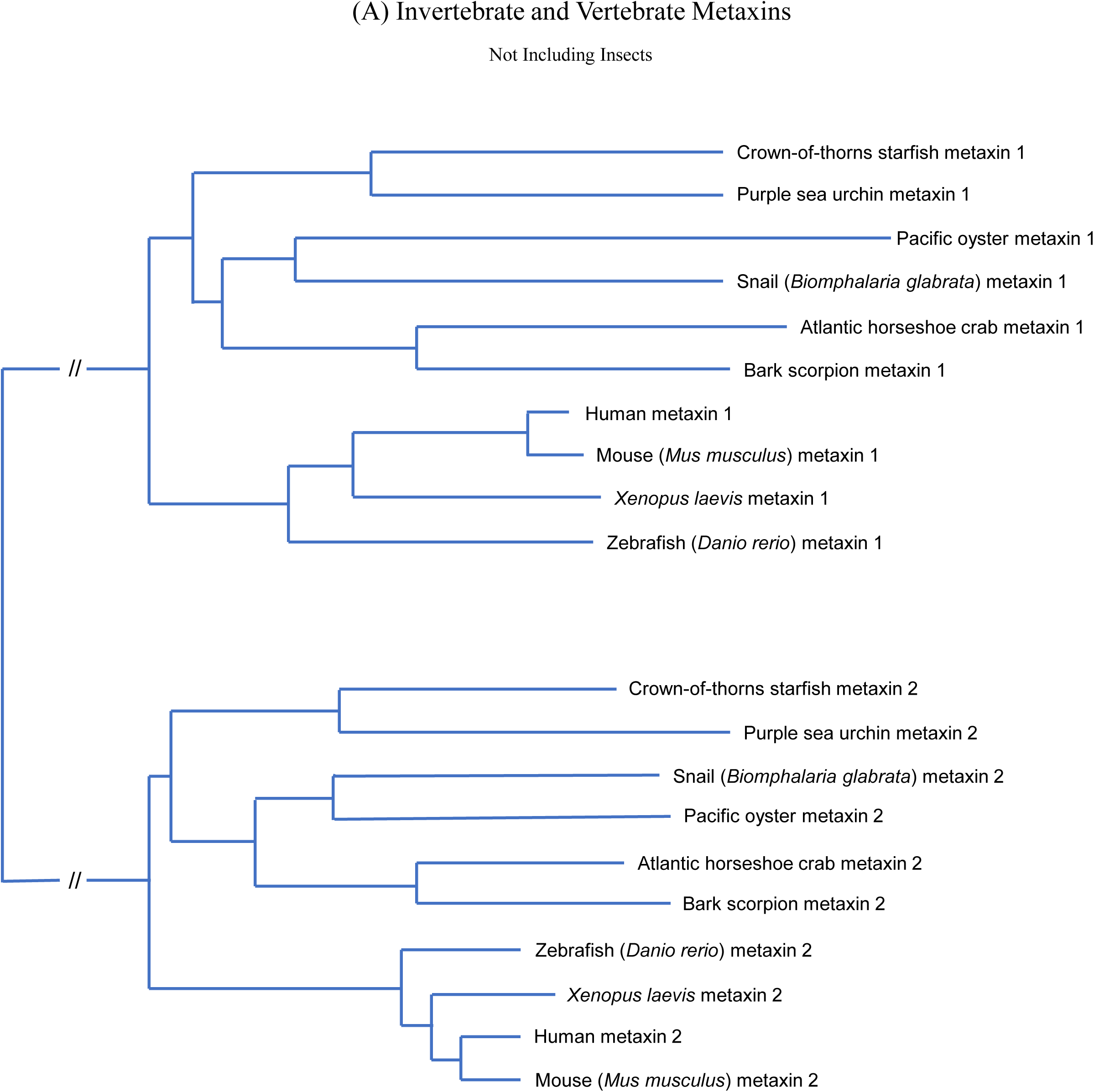
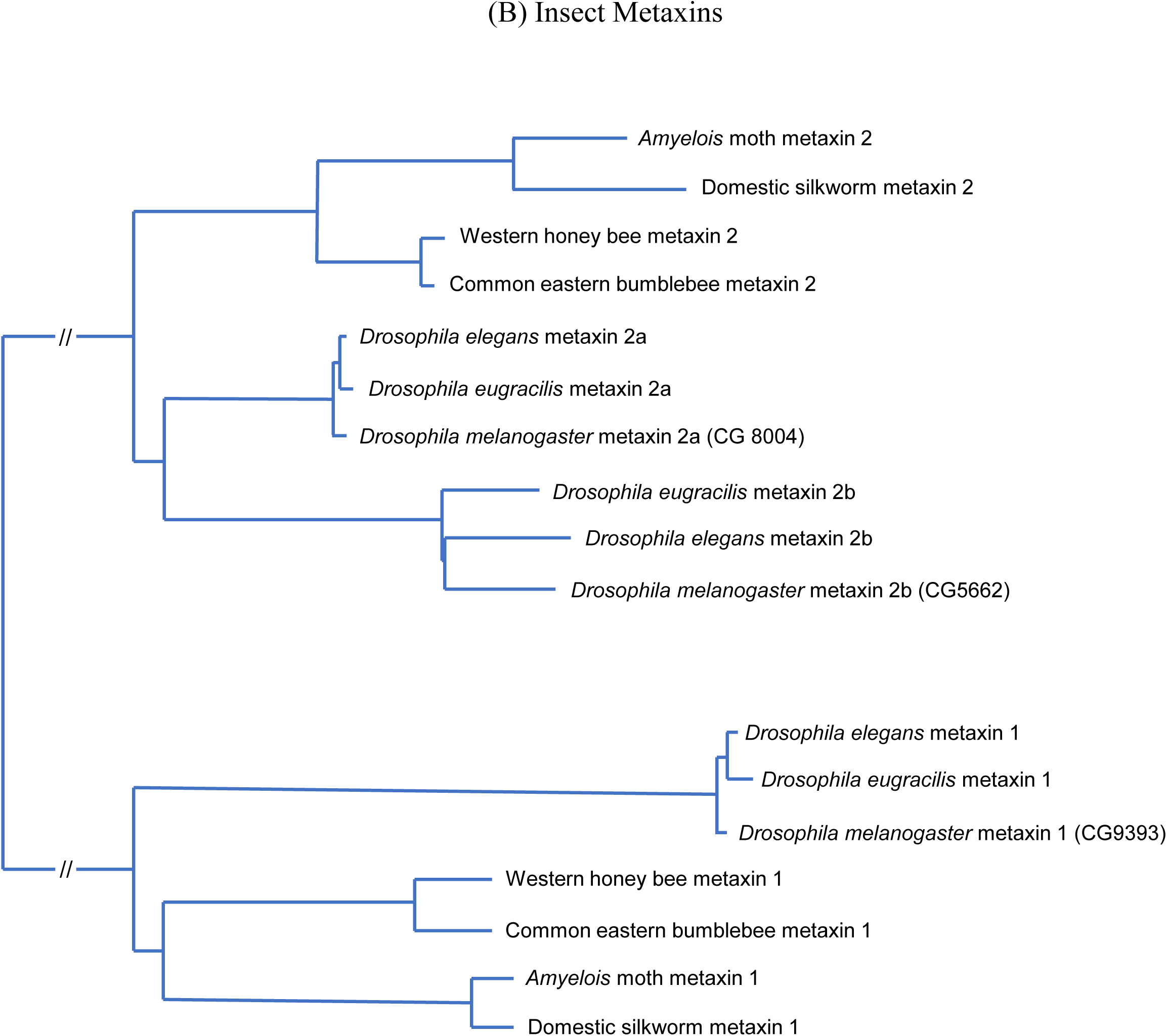
Evolutionary relationships of metaxin 1 and 2 proteins of invertebrates. (A) Phylogenetic tree of selected invertebrate and vertebrate metaxins. The invertebrate species shown are the crown-of-thorns starfish (*Acanthaster planci*), purple sea urchin *(Strongylocentrotus purpuratus*), Pacific oyster (*Crassostrea gigas*), snail (*Biomphalaria glabrata*), Atlantic horseshoe crab (*Limulus polyphemus*), and bark scorpion (*Centruroides sculpturatus*). The starfish and sea urchin are examples of the phylum Echinodermata, the oyster and snail are Mollusca, and the horseshoe crab and scorpion are Arthropoda. (B) The insect metaxins are those of a moth (*Amyelois transitella*), the domestic silkworm (*Bombyx mori*), the Western honey bee (*Apis mellifera*), and the common eastern bumblebee (*Bombus impatiens*). Three *Drosophila* species are also included (*Drosophila melanogaster, Drosophila elegans*, and *Drosophila eugracilis*). The moth and silkworm are examples of the taxonomic order Lepidoptera, the honey bee and bumblebee are Hymenoptera, and the *Drosophila* species are Diptera.

**Figure 4.**
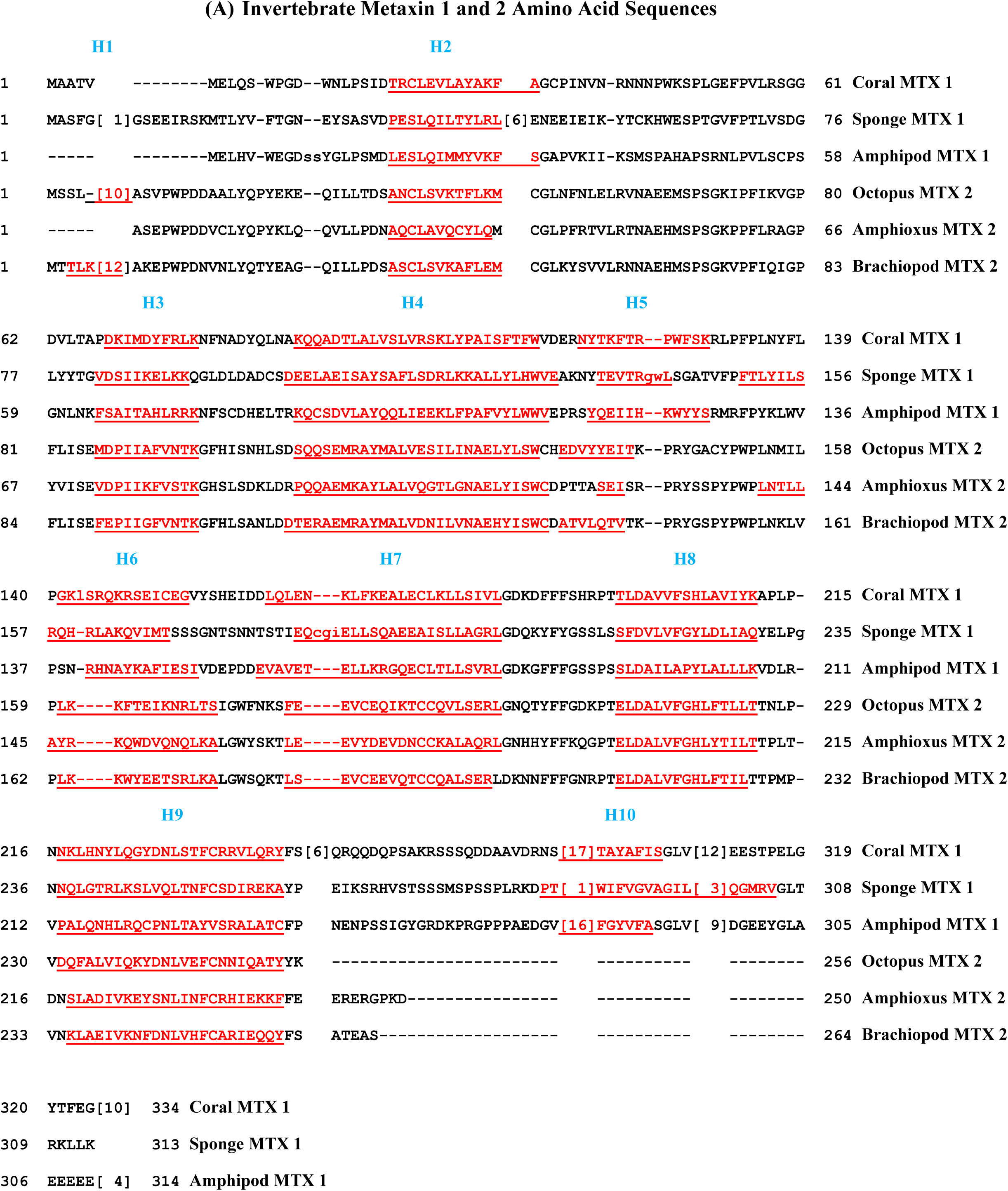
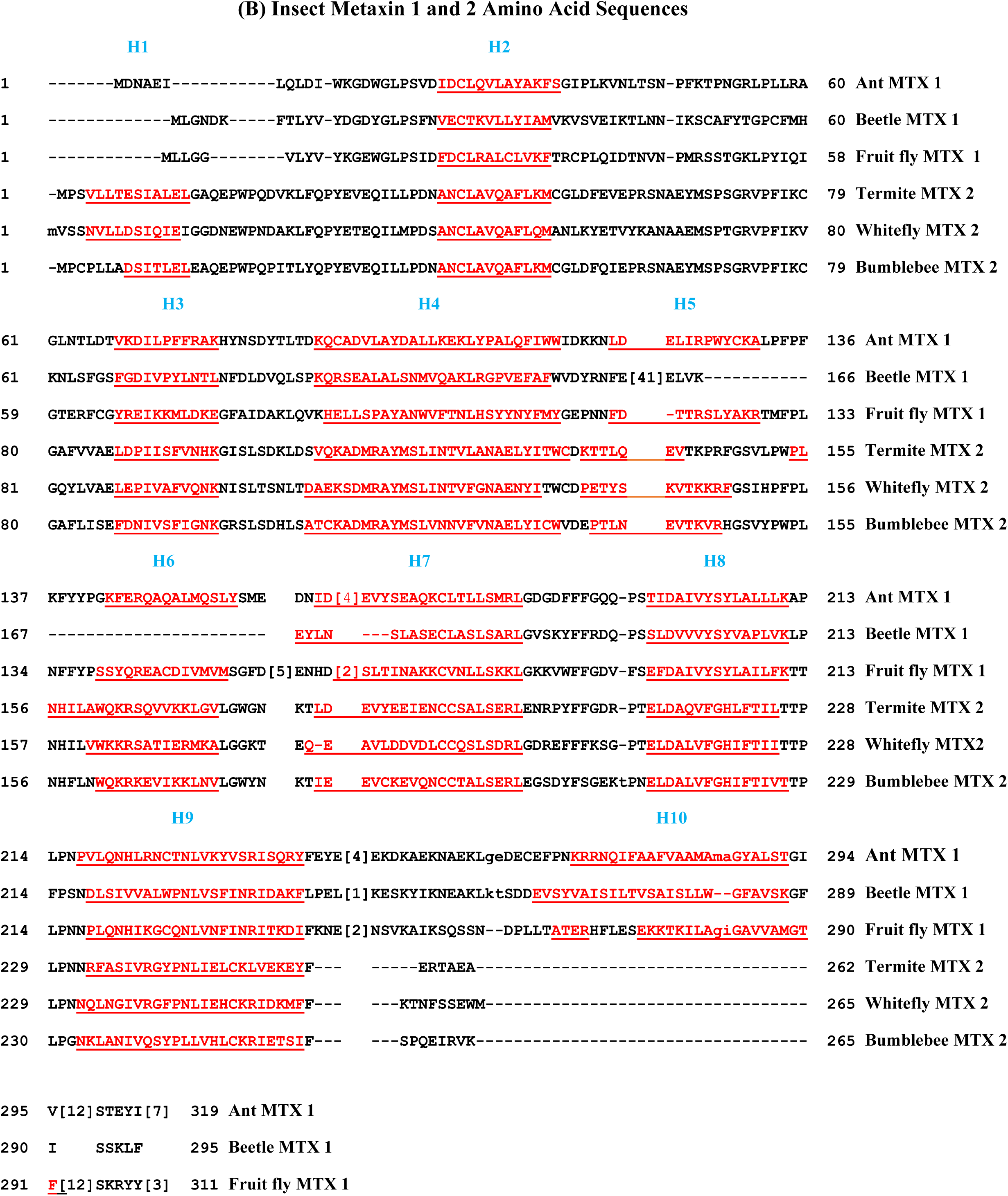
Secondary structures of invertebrate metaxin proteins: multiple sequence alignments. (A) Representative invertebrate metaxin 1 and metaxin 2 amino acid sequences are aligned, and the predicted alpha-helical segments are underlined and shown in red. The helices are labeled H1 through H10. The invertebrate species include a stony coral (*Acropora digitifera*; Phylum: Cnidaria), a sponge (*Amphimedon queenslandica*; Porifera), an amphipod crustacean (*Hyalella azteca*; Arthropoda), the California two-spot octopus (*Octopus bimaculoides*; Mollusca), the Florida lancelet or amphioxus (*Branchiostoma floridae*; Chordata), and a brachiopod (*Lingula anatina*; Brachiopoda). Invertebrate metaxin proteins, like the vertebrate metaxins, have predicted secondary structures that are dominated by alpha-helical segments, with very little beta-strand. The pattern of alpha-helical segments is highly conserved between species. Invertebrate metaxin 1 and metaxin 2 proteins each have 9 helices. Eight of the helices are shared by both metaxin proteins. Metaxin 1 has a C-terminal helix (H10) that metaxin 2 lacks, while metaxin 2 has an N-terminal helix (H1) that metaxin 1 lacks. The conservation of helices is striking, considering that the two metaxins only share about 20% sequence identities. For example, the metaxin 1 and 2 proteins of the California sea hare *Aplysia californica* (Mollusca) have only 21% identities. This result demonstrates how invertebrate metaxin secondary structure is more highly conserved than primary structure. (B) shows the alignment of representative insect metaxin 1 and 2 protein sequences. The alpha-helical segments, labeled H1 to H10, are underlined and in red. The insects included are the Florida carpenter ant (*Camponotus floridanus*; Order: Hymenoptera), the mountain pine beetle (*Dendroctonus ponderosae*; Coleoptera), the Mediterranean fruit fly (*Ceratitis capitata*; Diptera), the drywood termite (*Cryptotermes secundus*; Blattodea), the silverleaf whitefly (*Bemisia tabaci*; Hemiptera), and the common eastern bumblebee (*Bombus impatiens*; Hymenoptera). As in Figure 4(A), the insect metaxins have 9 alpha-helical segments that are highly conserved between species. Eight of the helices (H2 to H9) are found for both metaxin 1 and metaxin 2. Metaxin 2 has an extra N-terminal helix (H1), and metaxin 1 has an extra C-terminal helix (H10).

## 3. RESULTS AND DISCUSSION

### 3.1. Conservation of Metaxin Proteins Among Invertebrates

In this investigation, invertebrates were found to have distinct metaxin 1 and metaxin 2 genes like humans and other vertebrates. However, invertebrates were not observed to have metaxin 3 genes. Plants and bacteria are different and have a single metaxin-like gene (K.W. Adolph, unpublished). For invertebrates other than insects, metaxin 1 and 2 genes were found for representative invertebrates in a variety of phyla. These phyla include Echinodermata (example: sea urchin), Cnidaria (coral), Porifera (sponge), Chordata (amphioxus), Arthropoda (spider), Mollusca (octopus), Brachiopoda (brachiopod), Placozoa (*Tricoplax*), and Nematoda (*C. elegans*).

Insects were also found to have metaxin 1 and metaxin 2 genes, while metaxin 3 was not observed. This was true for representative insects (Phylum: Arthropoda; Class: Insecta) of various taxonomic orders: Diptera (example: fruit fly), Coleoptera (beetle), Lepidoptera (moth), Hymenoptera (honey bee), Hemiptera (bed bug), and Blattodea (termite).

For the fruit fly *Drosophila*, a second metaxin 2 gene was present in 13 of 17 *Drosophila* species studied. The two genes are designated metaxins 2a and 2b. The predicted 2a and 2b proteins have about 48% sequence identities. However, other insects have a single metaxin 2 gene, including the Mediterranean fruit fly, oriental fruit fly, and olive fruit fly. Different species of *Drosophila* have very similar metaxin proteins. For example, metaxin 1 proteins of different species have about 85% sequence identities, and metaxin 2a proteins 91%.

### 3.2. Amino Acid Sequence Homology of Invertebrate Metaxins

Invertebrate metaxin genes and proteins were first identified by homology with the amino acid sequences of human metaxin 1 and human metaxin 2 proteins. The invertebrate protein sequences are, in most cases, the sequences predicted from genomic DNA sequences. The identification of the invertebrate proteins as metaxins was strengthened by the presence of conserved, characteristic protein domains, the GST_N_Metaxin, GST_C_Metaxin, and Tom37 domains, but no additional major domains. Also, the lengths of the invertebrate metaxin proteins had to be comparable to human metaxin 1 (317 amino acids) or human metaxin 2 (263 amino acids).

The invertebrate metaxin 1 proteins share about 41% amino acid sequence identities with human metaxin 1. The metaxin 2 proteins share about 49% identities with human metaxin 2. As an example, the metaxin 1 protein sequence of the purple sea urchin *Strongylocentrotus purpuratus* (Echinodermata) has 46% identical amino acids when aligned with human metaxin 1. As additional examples, the sponge *Amphimedon queenslandica* (Porifera) has 28% identities, and the Pacific oyster *Crassostrea gigas* (Mollusca) has 39% identities. When aligned with the human metaxin 2 protein sequence, the sea urchin metaxin 2 sequence shows 50% identities, the sponge metaxin 2 sequence 35%, and the oyster sequence 47%.

Figure 1(A) includes the sequence of human metaxin 1 aligned with that of the roundworm *C. elegans* (*Caenorhabditis elegans*; Nematoda). There are 31% amino acid identities. In Figure 1(B), 36% identities are found in aligning human metaxin 2 with *C. elegans* metaxin 2. *C*.*elegans* is extensively used as a model organism for investigating the genetic control of development and physiology (Brenner, 1974; White et al., 1986), and was the first multicellular organism to have its genome completely sequenced (*C. elegans* sequencing consortium, 1998).

Figure 1(C) shows the alignment of human metaxin 1 with *Drosophila melanogaster* metaxin 1. The identities are 27%. For human and *Drosophila* metaxin 2a, 41% identities are observed (Figure 1(D)). The *Drosophila* metaxins are included because *Drosophila melanogaster* is widely used as a model organism for studies of molecular genetics, population genetics, and developmental biology (Adams et al., 2000; Wangler et al., 2017).

### 3.3. Protein Domains of Invertebrate Metaxins

A defining characteristic of metaxin proteins is the presence of GST_N_Metaxin and GST_C_Metaxin domains, and, in addition, Tom37 domains. Figure 2(A) shows the domain structure of representative invertebrate metaxin 1 and metaxin 2 proteins, while insects are in Figure 2(B). The invertebrates in 2(A) include the sea squirt *Ciona intestinalis*, which has the smallest genome of any chordate that can be utilized experimentally. As shown in the figure, *Ciona* metaxin 1 has GST_N_Metaxin1_like, GST_C_Metaxin1_3, and Tom37 domains. *Ciona* metaxin 2 has GST_N_Metaxin2, GST_C_Metaxin2, and Tom37 domains. The metaxin domains of the starlet sea anemone (*Nematostella vectensis*) and *Trichoplax adhaerens* are also included in Figure 2(A). The domains are identical to those of *Ciona*. The starlet sea anemone is a Cnidarian, the simplest animals with cells organized into tissues. *Trichoplax adhaerens* is the simplest known animal and has the smallest known genome of any animal. (For references, see Introduction.) The metaxin 1 and metaxin 2 domain structures of the three organisms are seen to be highly conserved.

The metaxin proteins of vertebrates, in particular humans, have similar domain structures to those seen in Figure 2. Human metaxin 1 and metaxin 3 have GST_N_Metaxin1_like, GST_C_Metaxin1_3, and Tom37 domains. Human metaxin 2 has GST_N_Metaxin2, GST_C_Metaxin2, and Tom37 domains.

The metaxin domains of representative insects are shown in Figure 2(B). The metaxin 1 and metaxin 2 domains are the same as those in 2(A) for invertebrates that don’t include insects. The examples in Figure 2(B) are the yellow fever mosquito (*Aedes aegypti*), a dampwood termite (*Zootermopsis nevadensis*), and the common bed bug (*Cimex lectularius*).

The Tom37 domain, as identified by the NCBI conserved domain database, is present in yeast (*Saccharomyces cerevisiae*) protein Tom37, now generally known as Sam37. Sam37 is a component of the SAM complex, the sorting and assembly machinery complex. The SAM complex, consisting of Sam50, Sam37, Sam35, and Mdm10, is located in the outer mitochondrial membrane, and is involved in the insertion of beta-barrel proteins into the outer membrane. The involvement of Sam37 (Tom37) in protein import into mitochondria of yeast is suggestive of a similar role for the metaxins in vertebrates and invertebrates, and such a role in human and mouse mitochondria has been experimentally demonstrated.

### 3.4. Evolutionary Relationships of Invertebrate Metaxin Proteins

Phylogenetic analysis demonstrates that the invertebrate metaxin 1 and metaxin 2 homologs are grouped separately from vertebrate metaxins, including human metaxins. Figure 3(A) shows the phylogenetic relationships of selected invertebrate and vertebrate metaxins. The analysis was based on multiple sequence alignments of the protein sequences. The horizontal lengths of the branches are proportional to the extent of evolutionary change. As Figure 3(A) demonstrates, the metaxin 1 proteins form a distinct group relative to the metaxin 2 proteins. And within each group, the invertebrate and vertebrate proteins form separate clusters.

Also within each group, invertebrate metaxins of the same phylum are seen to be the most closely related. This includes the starfish and sea urchin metaxin 1 and metaxin 2 proteins (Echinodermata), the oyster and snail metaxins (Mollusca), and the horseshoe crab and scorpion metaxins (Arthropoda). Similarly, for vertebrate metaxin 1 and metaxin 2 proteins, the human and mouse proteins are more closely related compared to the *Xenopus* and zebrafish metaxins.

The observed grouping of the invertebrate and vertebrate metaxin 1 proteins indicates that the metaxin 1 proteins arose from a common ancestor. The separate grouping of the metaxin 2 proteins likewise indicates that the metaxin 2 proteins originated from a common ancestor. In addition, the results strengthen the likelihood of a common ancestral gene sequence for both of the metaxins. This is true even though alignments of the metaxin 1 and 2 protein sequences show relatively low percentages of amino acid identities (22% for human metaxins 1 and 2).

Insect metaxins are included in Figure 3(B). As with the other invertebrate metaxins in Figure 3(A), the insect metaxin 1 and metaxin 2 proteins form separate groups. The metaxins of insects of the same taxonomic order are seen to be more closely related than those in different orders. The examples shown are moth and silkworm metaxins (Order: Lepidoptera), honey bee and bumblebee metaxins (Hymenoptera), and *Drosophila* metaxins of different species (Diptera). Although two distinct groupings are found for the metaxin 1 and metaxin 2 proteins, the insect metaxins, like other invertebrate metaxins, most likely originated from an ancestral metaxin, even though the amino acid sequences of metaxins 1 and 2 have diverged significantly.

The metaxins of the three species of *Drosophila* in Figure 3(B) are very closely related phylogenetically. For metaxin 2, the two *Drosophila* proteins (metaxins 2a and 2b) found for a number of *Drosophila* species are divided into a metaxin 2a cluster and a metaxin 2b cluster. The results indicate that the 2a proteins are related through a common ancestor, and similarly for the 2b cluster. And both the 2a and 2b proteins are likely to have originated from a common metaxin 2 ancestral sequence.

### 3.5. Neighboring Genes of Invertebrate Metaxins

The genes adjacent to invertebrate metaxin 1 and metaxin 2 genes are largely different comparing one invertebrate with another. Also, no similarities are seen with the arrangement of genes in humans and other vertebrates, where the metaxin 1 gene (*MTX1*) is next to the thrombospondin 3 gene (*THBS3*), the metaxin 2 gene (*MTX2*) is adjacent to a cluster of homeobox genes, and the metaxin 3 gene (*MTX3*) is next to the thrombospondin 4 gene (*THBS4*). In contrast, invertebrate metaxin genes are not adjacent to thrombospondin or homeobox genes. And the genes in the chromosomal regions up to 10 to 20 or more genes from the metaxin genes are different than the genes of humans and other vertebrates, and generally differ from one invertebrate to another. For example, the roundworm *C. elegans*, which has a well-annotated genome, has a metaxin 1 gene (*mtx-1*) between the humpback gene and a heat-shock protein gene. And a metaxin 2 gene (*mtx-2*) is next to the prolyl carboxy peptidase gene. Unique arrangements of genes adjacent to the metaxin genes were also found for *Ciona intestinalis*, a sea squirt, with the metaxin 1 gene (*mtx-1*) between genes for a regulatory protein and a transforming, acidic coiled-coil protein. There are exceptions, however, to having different neighboring genes. A few genes were found to be close to the metaxin genes of some, but not all, invertebrates. As an example, an ADP-dependent glucokinase gene is close to 3 of 10 invertebrate metaxin 1 genes. But the metaxin 2 genes of the 10 invertebrates showed no similarities in adjacent genes.

More specifically for insects, the genes near metaxin 1 genes are different for *Drosophila melanogaster* compared to the Mediterranean fruit fly, honey bee, etc. Also, for *Drosophila melanogaster*, the genes close to the two metaxin 2 genes (2a and 2b) are different. However, the genes adjacent to the single metaxin 1 homolog are conserved in ten of twelve *Drosophila* species. Other exceptions to having different genes are closely related species such as the oriental fruit fly and olive fruit fly, which have the same 5 adjacent genes.

### 3.6. Secondary Structures of Invertebrate Metaxin Proteins

Alpha-helical segments are the dominant secondary structures of invertebrate metaxins. There is relatively little beta-strand. Figure 4 shows that representative invertebrate metaxin 1 and 2 proteins each have eight predicted helical segments (H2 through H9) with the same pattern of spacing between the segments. Metaxin 1 proteins have an additional 9th helical segment (H10) in their extended C-terminal regions, while metaxin 2 proteins have a 9th helix (H1) in their N-terminal regions. This pattern of alpha-helices is the same as the pattern for metaxins 1, 2, and 3 of humans and other vertebrates. As shown in Figure 4(A), the conserved pattern of helices was found for invertebrate phyla including Cnidaria (example: coral), Porifera (sponge), Arthropoda (amphipod), Mollusca (octopus), Chordata (amphioxus), and Brachiopoda (brachiopod). Figure 4(B) includes multiple sequence alignments showing the pattern of conserved helices for a selection of insects (Phylum: Arthropoda; Class: Insecta). The insect taxonomic orders represented in Figure 4(B) are Hymenoptera (ant, bumblebee), Coleoptera (beetle), Diptera (fruit fly), Blattodea (termite), and Hemiptera (whitefly). The pattern of predicted alpha-helical segments is very similar to the pattern in Figure 4(A) for representative invertebrates, not including insects. Eight helices (H2 to H9) are found for both metaxin 1 and metaxin 2 of insects. H1, an extra N-terminal helix, is characteristic of metaxin 2 proteins. H10, an additional C-terminal helix, characterizes metaxin 1 proteins. These highly conserved patterns are found for other insects and non-insect invertebrates not included in Figure 4.

Metaxin 1 proteins of vertebrates have a predicted transmembrane helix near the C-terminus. However, vertebrate metaxin 2 and metaxin 3 proteins don’t have a C-terminal transmembrane helix. The metaxin 1 transmembrane helix is identified as helix H10 of vertebrates, since it largely coincides with H10. The transmembrane helix could serve to anchor metaxin 1 proteins to the outer mitochondrial membrane in connection with their role in protein import into mitochondria. The presence of a C-terminal transmembrane helix is also predicted for many invertebrate metaxin 1 proteins. For example, the sea urchin *Strongylocentrotus purpuratus* has a transmembrane helix between residues 272 and 294 of the 319 amino acid protein. Belcher’s lancelet or amphioxus metaxin 1 has the helix between residues 269 and 291 of the 321 amino acid protein. The transmembrane helix mostly coincides with C-terminal helix H10. But neither of the metaxin 2 proteins of these invertebrates has a transmembrane helix. Considering insects, the *Drosophila melanogaster* metaxin 1 homolog (CG9393) has a transmembrane helix between residues 281 and 301 of the 316 amino acid protein. But neither of the two *D. melanogaster* metaxin 2 homologs (CG5662 and CG8004) has a transmembrane helix.

